# Pick-ya actin: A method to purify actin isoforms with bespoke key post-translational modifications

**DOI:** 10.1101/833152

**Authors:** Tomoyuki Hatano, Lavanya Sivashanmugam, Andrejus Suchenko, Hamdi Hussain, Mohan K. Balasubramanian

**Affiliations:** Centre for Mechanochemical Cell Biology and Division of Biomedical Sciences, Warwick Medical School, University of Warwick, Coventry CV4 7AL, United Kingdom

## Abstract

Actin is one of the most abundant eukaryotic cytoskeletal polymer-forming proteins, which in the filamentous form regulates a number of physiological processes, ranging from cell division and migration to development and tissue function. Actins are differentially post-translationally modified (PTMs) in different organisms, which include Met, Ala, Asp, and Glu *N*-acetylation, *N*-arginylation, and the 73th His residue (His-73) methylation, with different organisms displaying a distinct signature of PTMs. Currently methods are not available to produce actin isoforms with organism specific PTM profile. Here we report Pick-ya actin, a method to express actin isoforms from any eukaryote with its own key characteristic PTM pattern. We achieve this using a synthetic biology strategy in a yeast strain that expresses 1. actin isoforms with the desired *N*-end via ubiquitin fusion and 2. mammalian enzymes that promote acetylation and methylation. Pick-ya actin should greatly facilitate biochemical, structural, and physiological studies of the actin cytoskeleton and its PTMs.

## Introduction

Actin is a highly conserved eukaryotic cytoskeletal protein, which polymerizes into double stranded, dynamic polar filaments that exhibit continuous polymerization and depolymerization. A variety of eukaryotes have evolved diverged actin isoforms and orthologues, which assemble into a variety of molecular machineries executing mechanical tasks including cell division and cell migration. How actins undergo defined molecular structural changes facilitating polymerization and depolymerization and how they carry out distinct functions at different cellular locations and in different organisms and tissues is not well understood. The understanding of actin function in different organisms and tissues is further complicated by the vast array of PTMs, which differ among different organisms. Some of the known modifications of actin include 1. Met *N*-acetylation in yeast; 2. Ala *N*-acetylation in *Arabidopsis thaliana* (Bienvenut et al., 2012); 3. Asp or Glu *N*-acetylation in animals and amoeba, mediated by *N*-acetyl transferase NAA80 (Drazic et al., 2018; Goris et al., 2018); 4. *N*-arginylation in mammals (Karakozova et al., 2006); and 5. His-73 methylation in animals and amoeba, mediated by the His methyl transferase SETD3 (Kwiatkowski et al., 2018; Wilkinson et al., 2018). Thus, precise biochemical and biophysical understanding of actin function in various organisms and tissues depends on the ability to produce actin isoforms with organism-specific signature PTMs. The current methods for actin purification either constitutively introduce Asp/Glu *N*-acetylation and His-73 methylation (purification from rabbit or chicken muscle, Sf9-insect cell cultures, *Dictyostelium*, and human platelets) or do not introduce either of the above-mentioned modifications (*Saccharomyces cerevisiae* and *Pichia pastoris*) (Hatano et al., 2018; Müller et al., 2013; Noguchi et al., 2007; Takashi et al., 2009).

Given that recombinant actin expressed in *P.pastoris* is neither Asp/Glu *N*-acetylated nor His-73 methylated, we reasoned that further engineering of *P.pastoris* may provide a means to express and purify actin isoforms with distinct organism-specific PTMs. Here we express NAA80 and SETD3 either singly or together in *P.pastoris*. This combined with our previously described ubiquitin-fusion approach (Hatano et al., 2018) has led to the development of Pick-ya actin, which facilitates production of actin isoforms with organism specific PTM profiles.

## Results and Discussion

Figure 1 shows key PTMs observed in actin from mammals, yeast, and the plant *A. thaliana*. Mammalian actin can be isolated with one of seven different modifications at the *N*-terminus, which are mediated by a combination of aminopeptidases, NAA80 acetyl transferase (Drazic et al., 2018; Goris et al., 2018), and ATE1 arginyl-tRNA-protein transferase (Karakozova et al., 2006). Furthermore, human actin is methylated on His-73 by the SETD3 methyl transferase (Kwiatkowski et al., 2018; Wilkinson et al., 2018). *A. thaliana* actin is *N*-terminally acetylated at alanine and is only weakly (9%) methylated on His-74, equivalent to mammalian and yeast His-73 (Wilkinson et al., 2018). Finally yeast actin is acetylated on the initiator Met and is not methylated on His-73 (Kalhor et al., 1999). We systematically established yeast strains that introduced one or more of the above-mentioned modifications. Towards this end, we cloned human NAA80 acetyl transferase and human SETD3 methyl transferase in *P. pastoris* expression plasmids, which are described in the subsequent sections.

**Figure 1.**
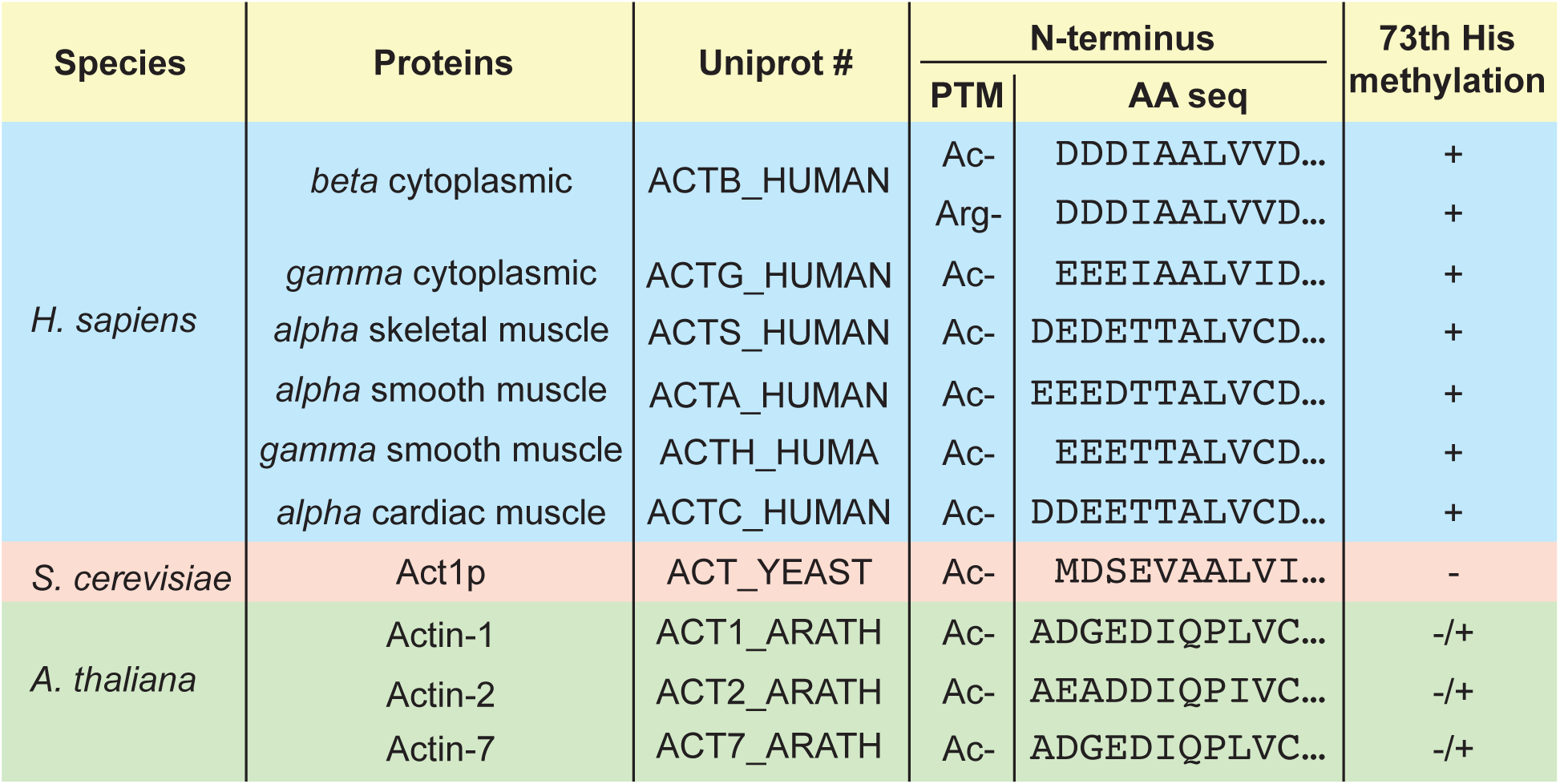
Post-translational modification of actin. Divergent post-translational modifications on actin. Shown are *N*-terminal peptide sequences of nine actins (6 from *H. sapiens*, 1 from *S. cerevisiae*, and 3 from *A. thaliana*) and the observed modifications at the *N*-terminus. The right column summarizes if His-73 methylation is observed in these organisms.

### Expression in *P. pastoris* and purification of *N*-acetylated human β and γ actin

Human β and γ actin expressed in *P.pastoris* are not processed at the *N*-terminus, as a result methionine is retained at the *N*-terminus and is acetylated by native acetyl transferases (Hatano et al., 2018). The canonical *N*-acetylation occurs on Asp residue on β actin and Glu residue on γ actin. To facilitate generation of β and γ actin with Asp or Glu at the *N*-terminus, we expressed a DNA molecule (Figure 2A) encoding a translational fusion of ubiquitin with the actin gene lacking the initiator methionine followed by sequences encoding human thymosin β4 (Tβ4) and the 8 consecutive His amino acid residues (8His), which facilitates one-step affinity purification using nickel-NTA resin (Hatano et al., 2018; Huang et al., 2016; Kijima et al., 2016; Noguchi et al., 2007). In this strain, we introduced an integrative plasmid vector backbone or a plasmid expressing human NAA80, which promotes actin *N*-acetylation. The translational fusion was purified as described previously using an affinity-binding approach (Hatano et al., 2018). When the four purified actin-fusions were subjected to mass spectrometry, we found that the majority of *N*-terminus in β actin contained an Asp residue, whereas a large number of Ac-Asp peptides were detected when NAA80 was co-expressed in these cells (Figure 2B and D). Similarly, majority of γ actin contained a Glu residue at its *N*-terminus in control cells, whereas it contained Ac-Glu at the *N*-terminus when NAA80 was co-expressed (Figure 2C and E). Evaluation of the number of non-acetylated vs acetylated *N*-terminal peptides from β and γ actin clearly demonstrated that NAA80 expression strongly promotes the *N*-terminal acetylation of β and γ actin. Together, we concluded that expression of NAA80 was sufficient to introduce *N*-acetylation of human β and γ actin expressed in *P. pastoris*.

**Figure 2.**
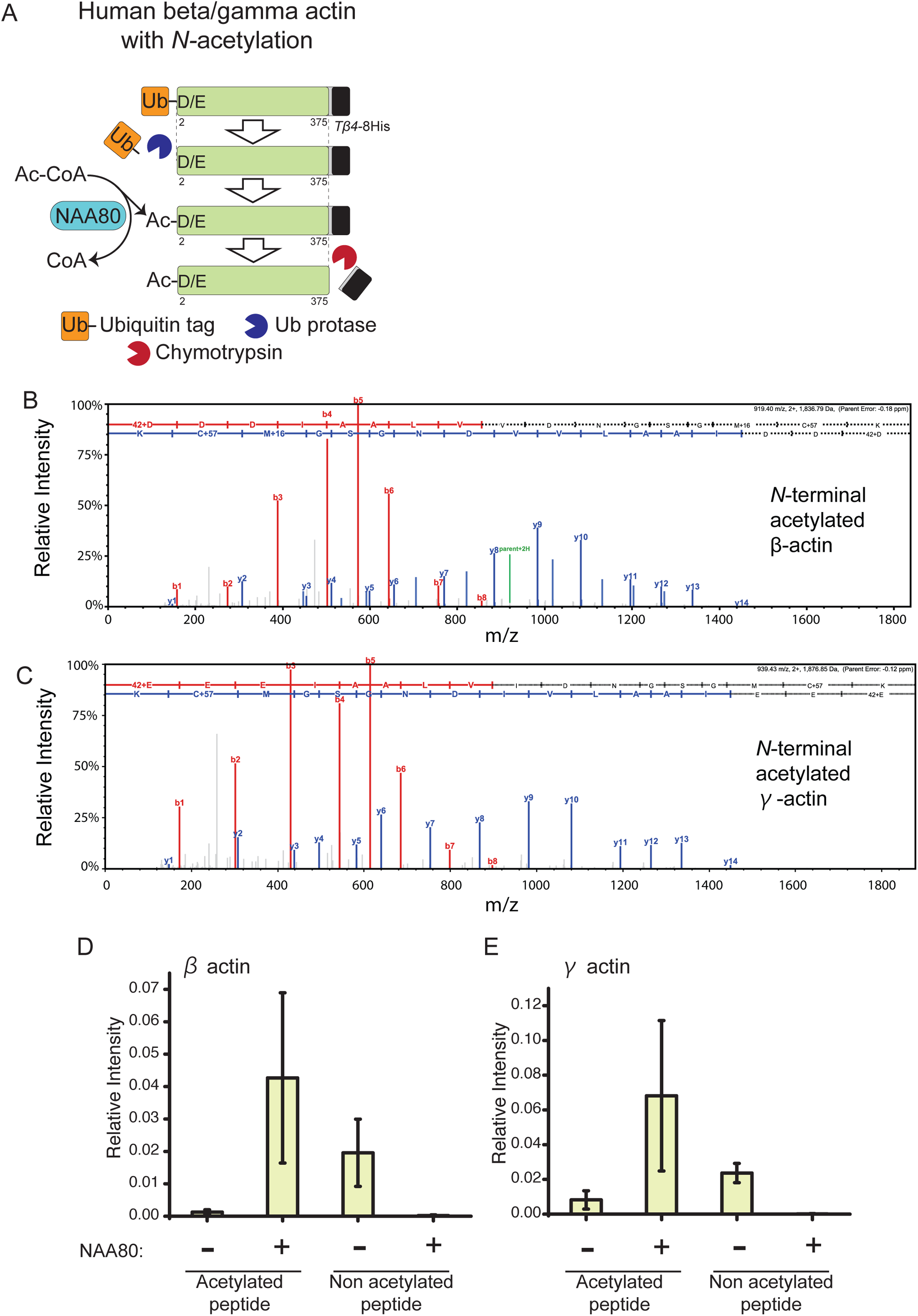
*N*-acetylation of recombinant human β and γ actin. A. Schematic representation of purification of synthetically *N*-acetylated human β and γ actin. A Ubiquitin (Ub)-moiety was fused at the *N*-terminus of actin lacking the first Met. The Ub-tag is removed by the endogenous Ub protease and the exposed *N*-terminal Asp in β actin or Glu in γ actin is acetylated by the NAA80 enzyme. The *C*-terminal tag is then removed by chymotrypsin treatment to liberate modified full length actin. B and C. MS spectra of the *N*-terminal peptide derived from recombinant β actin (B) and γ actin (C). D and E. Label-free quantification of acetylated or unmodified *N*-terminal peptide in β and γ actin. The bar-graph shows mean +/-SD from three independent experiments. The relative intensity of *N*-acetylated peptide in control sample and the sample expressing NAA80 is shown. (D) β actin, (E) γ actin.

### Expression in *P. pastoris* and purification of His-73 methylated human β and γ actin

We next established a system to prepare His-73-methylated human β and γ actin. Recent work has shown that the mammalian SETD3 methyl-transferase methylates His-73 of human actins, using *S*-adenosyl methionine as a methyl group donor (Kwiatkowski et al., 2018; Wilkinson et al., 2018; Figure 3A). To promote actin His-73 methylation, we co-expressed the mammalian SETD3 and β or γ actin in *P.pastoris* cells. Again, methylated peptides covering His-73 were detected in cells expressing SETD3, but not in cells without SETD3 (Figure 3B-E). Both, β and γ actin were methylated on His-73 (Figue 3B and C). Quantitation of relative intensity of methylated vs non-methylated peptides revealed a clear correlation between expression of SETD3 and methylation of His-73 on β and γ actin expressed in *P. pastoris* (Figure 3D and E). These experiments established that human β and γ actins can be efficiently methylated by SETD3 in *P. pastoris*.

**Figure 3.**
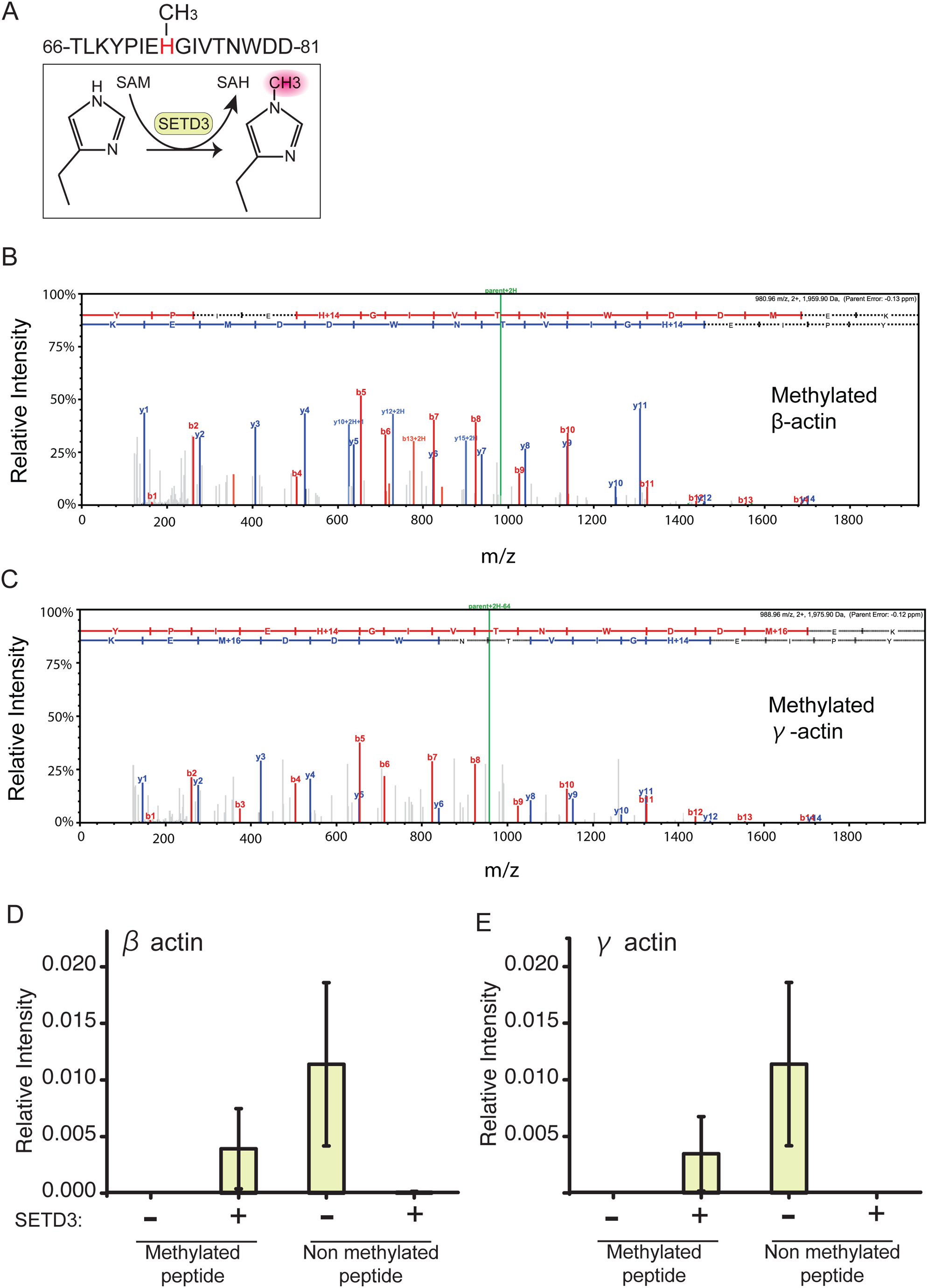
Methylation of the 73th His residue in human β and γ actin. A. Enzymatic reaction of SETD3 methyl transferase. Peptide sequence from human non-muscle actin around the 66-81th amino acid residues are shown. SETD3 recognizes the peptide surrounding His-73 in actin and transfers a methyl group from *S*-adenosyl methionine (SAM) to the His residue to generate methylated actin and *S*-adenosyl homocysteine (SAH). B and C. MS spectra of the peptide from recombinant actin carrying methylation at His-73. (B) β actin, (C) γ actin. D and E. Label-free quantification of methylated or unmodified His-73 Peptide in β and γ actin. The bar-graph shows mean +/-SD from three independent experiments. The relative intensity of methylated peptide in control sample and the sample expressing SETD3 is shown. (D) β actin, (E) γ actin.

### Expression in *P. pastoris* and purification of *N*-acetylated and His-73 methylated human β actin and arginylated and His-73 methylated actin

Next, we sought to purify dually modified human β actin with *N*-acetylation and His-73 methylation. Towards this goal, we employed a ubiquitin-fusion strategy as described in Figure 2A. In *P. pastoris* cells expressing the Ub-D-actin-Tβ4-8His fusion, we expressed NAA80 and SETD3. Actin was purified sequentially by nickel affinity chromatography, release from Tβ4 and 8His by chymotrypsin treatment, and a cycle of polymerization and depolymerization (see materials and methods for full details of the purification). Figure 4A shows results from a size exclusion column, in which monomeric actin was obtained in the peak fractions. Mass spectrometric analysis of the purified actin confirmed that it was both *N*-acetylated and His-73 methylated (Supplemental figure 3), as described in Figures 2 and 3.

**Figure 4.**
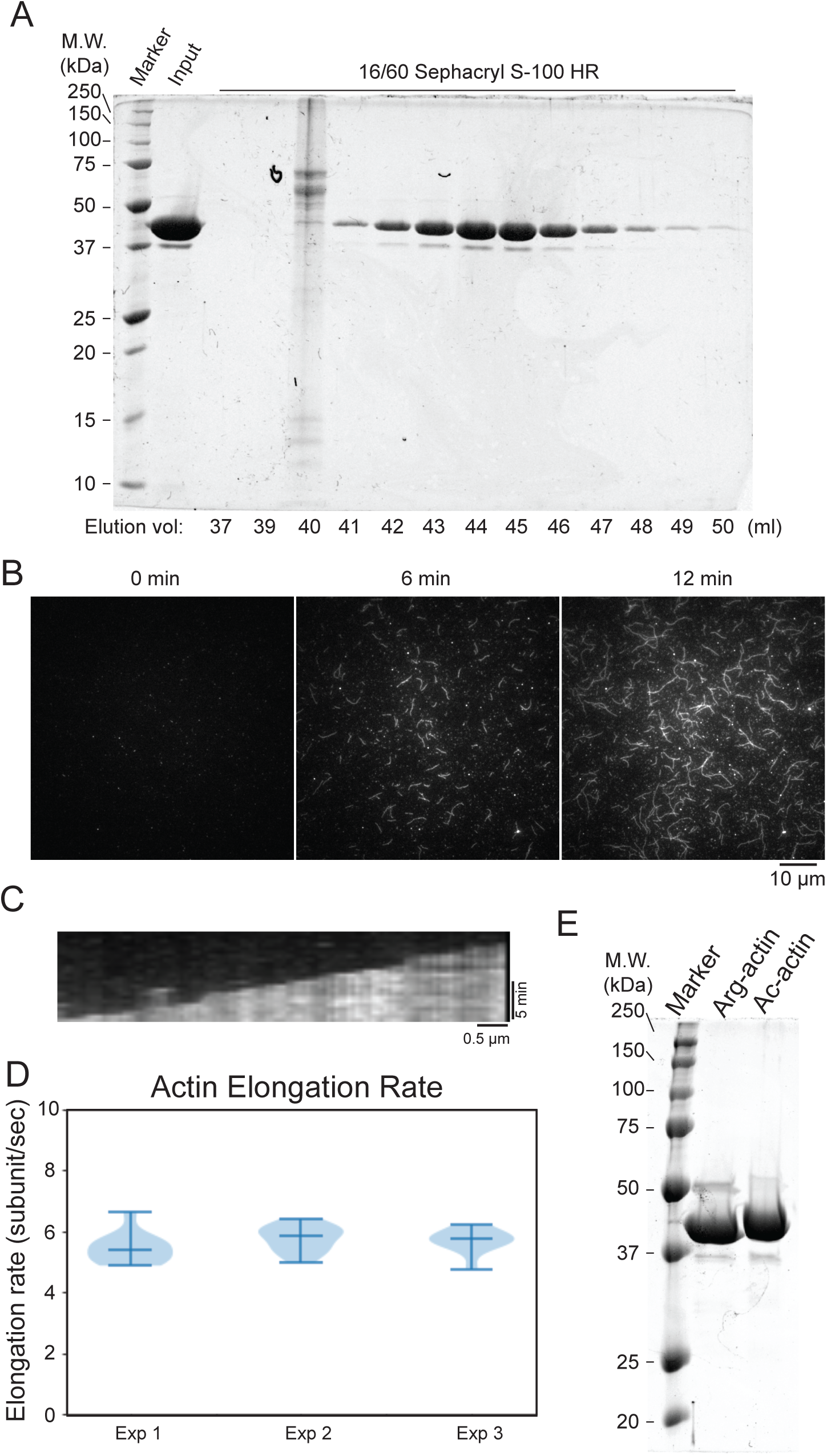
Purification of dually modified human β-actin. A. Size-exclusion chromatography (Sephacryl S-100HR) showing homogenous G-actin purified from *P. pastoris* expressing NAA80 and SETD3. B. Time lapse images showing actin polymerization. Actin was visualized with 20% Sc Act1 with Alexa Fluor 488 and tethered on the coverslip surface with 0.5% ScAct1 with biotin-PEG2. C. Kymograph (right) showing elongation of a single actin filament. D. Elongation rate of actin. Elongation rate at the barbed end of actin was quantified as described in materials and methods. E. SDS-PAGE of purified arginylated β actin carrying methylation at His-73. *N*-acetylated and His-73 methylated β-actin is also shown in this gel.

Next, we tested if the purified *N*-acetylated and His-73 methylated human β actin was capable of polymerization. Towards this goal, we incubated the purified human actin with a trace amount of *S. cerevisiae* actin that was fluorescently labeled by conjugation of Alexa Fluor 488 C5-maleimide to Cysteine 374. In this experiment, we found that the dually modified actin was capable of polymerization with an elongation rate of ∼ 6 subunits / second in three independent experiments (Figure 4B-D).

While *N*-acetylation is one key modification of the *N*-terminus of actin, a mutually exclusive modification termed arginylation also occurs on human β actin. We, and others, have shown that the ubiquitin fusion strategy can be used to express and purify arginylated human β actin from *P. pastoris*. To develop a full actin toolkit, we sought to express and purify arginylated and methylated actin. To this end, we co-expressed Ub-R-actin-Tβ4-8His fusion and SETD3 in *P.pastoris*. We were able to purify arginylated and His-73 methylated actin at levels comparable to *N*-acetylated and His-73 methylated actin (Figure 4E).

Collectively, these experiments showed that the system we have developed can allow a mix and match strategy to purify actin in 1. unmodified, 2. *N*-actelayed, unmethylated 3. *N*-acetylated, methylated 4. arginylated, unmethylated, and 5. arginylated and unmethylated forms.

### Expression in *P. pastoris* and purification of Ala *N*-acetylated and His-73 methylated *A. thaliana* actin

We next wanted to extend the utility of the approach we have developed to purify actin isoforms. We were in particular interested to express plant actin isoforms since they have unique profiles of PTMs (Figure 1). Actins from the model plant *A. thaliana* have been shown through proteomic analysis to be *N*-acetylated on alanine (rather than Asp or Glu, as seen in mammalian actins) and methylation of His-74 is only found in ∼9% of the cellular pool of actin (Bienvenut et al., 2012; Wilkinson et al., 2018). We reasoned that the *P. pastoris* expression system might offer a better avenue to producing plant actins due to our ability to control each of the modifications. We generated a fusion *A. thaliana* Actin-2 (Act2) gene, in which we again used the ubiquitin fusion strategy, to allow exposure of the second Ala residue at the *N*-terminus (Figure 5A). A Ub-A-actin-Tβ4-8His fusion was expressed in *P.pastoris*. We found through mass spectrometry that the exposed *N*-terminal alanine residue was acetylated by a native acetyl transferase (Figure 5B). This observation established that *A. thaliana* actin can be expressed in *P. pastoris* in a *N*-acetylated form for biochemical studies.

**Figure 5.**
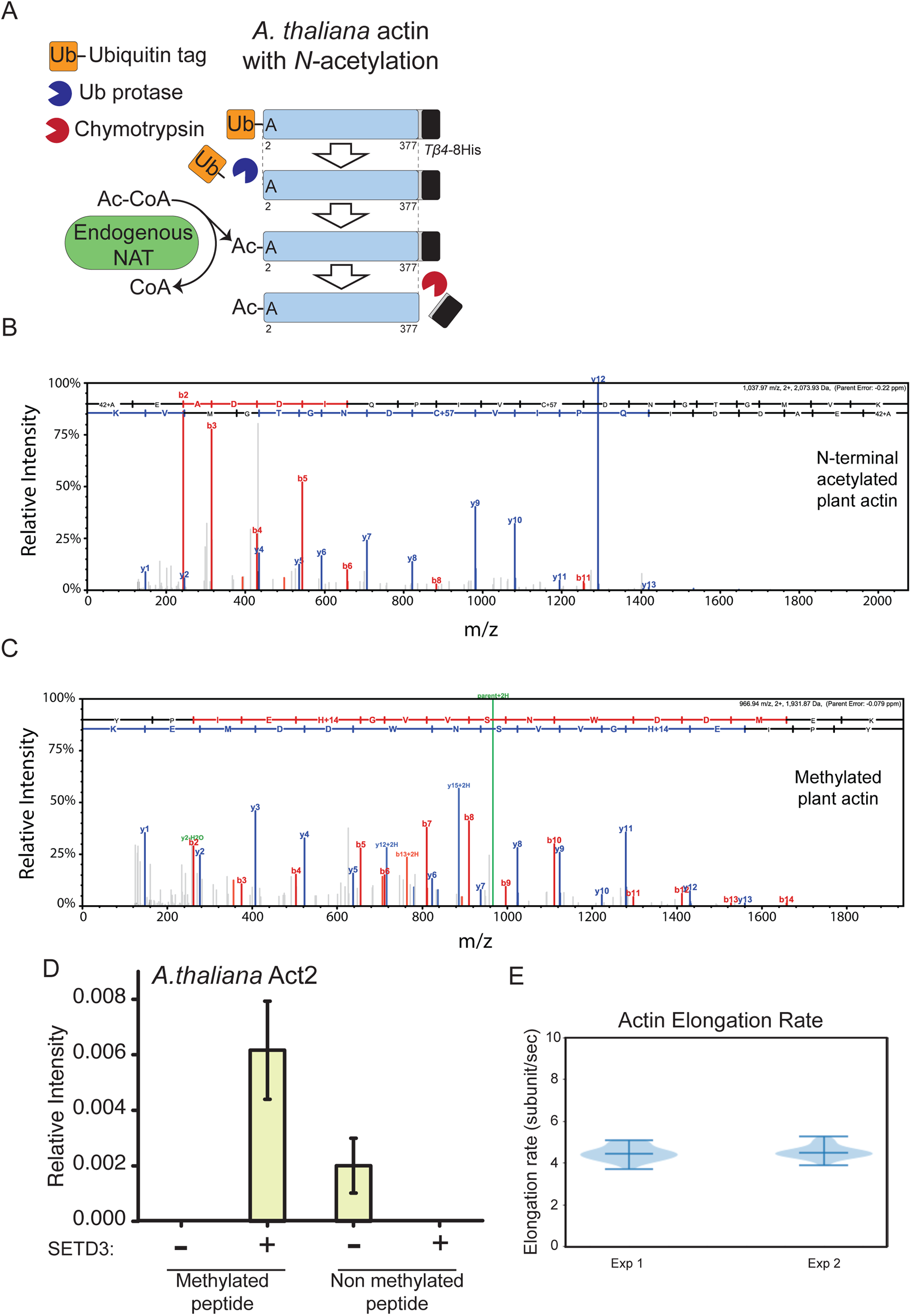
*A. thaliana* Actin-2 with appropriate PTMs. A. Schematic representation of purification of *A. thaliana* actin. A Ub-moiety was fused at the *N*-terminus of actin lacking the first Met. The Ub-tag is removed by the endogenous Ub protease and the exposed *N*-terminal Ala is *N*-acetylated by a native acetyl transferase. The *C*-terminal tag is then removed by chymotrypsin treatment to liberate modified full length actin. B. MS spectra of acetylated peptides in recombinant *A.thaliana* Actin-2. C. Methylation at His-74 in *A. thaliana* Actin-2 in the presence of SETD3. D. Label-free Quantification of methylated and unmodified His-73 Peptide in Arabidopsis thaliana (At) actin. The bar-graph shows mean +/-SD from two independent experiments. The relative intensity of methylated peptide in control sample and the sample expressing SETD3 is shown. E. Elongation rate of recombinant *A. thaliana* Actin-2. Elongation rate at the barbed end of actin was quantified as described in materials and methods.

Since ∼ 9% of *A. thaliana* actin is methylated on His-74 (corresponds to His-73 in animal actin), we desired to establish a system to purify His-74 methylated actin (Wilkinson et al., 2018). To this end, we expressed SETD3 in *P.pastoris* expressing Ub-A-actin-Tβ4-8His. Mass spectrometry confirmed that the *A. thaliana* actin was methylated on His-74 (Figure 5C and D). Comparison of the relative intensity profiles of methylated and non-methylated peptides revealed a clear correlation between SETD3 expression and His-74 methylation (Figure 5D). This work established that *A.thaliana* actin can be expressed in an alanine *N*-acetylated form with or without His-74 methylation.

Finally, we tested if the purified Ala N-acetylated *A. thaliana* actin was capable of polymerization *in vitro*. We performed a similar experiment to that with the human β-actin described in Figure 4B-D, and found that *A. thaliana* actin polymerized at ∼ 4 monomers / second in two independent experiments (Figure 5E). This finding established that A. thaliana actin purified from *P. pastoris* was functional *in vitro*.

## Conclusions

In this study, we have substantially improved upon our previous work (Hatano et. al., 2018) on expression and purification of heterologous actin isoforms in *P.pastoris*. Importantly, we have developed approaches to precisely control the *N*-acetylation and His-73 methylation status of the expressed heterologous actins. We see two major advantages for this system we describe, which we term as Pick-ya actin.

First, Pick-ya actin will facilitate biochemical and structural studies of actin isoforms, and their modifications. We imagine that any laboratory investigating the role of each of the modifications of mammalian β and γ actin on its structural, biochemical, and biophysical properties may be able to work with pure actin isoforms with the desired modifications. Actin without modifications can be readily isolated without the need for cell-lines lacking NAA80 or SETD3 and actins with modifications can be purified upon co-expression of the appropriate enzymes.

Second, and perhaps more importantly, Pick-ya actin will help understand the diversity of actin function in a variety of different organisms with different profiles of *N*-acetylation and His-73 methylation. In this regard, Pick-ya actin may be preferable to the Sf9 insect cell culture-based approach. This is due to the fact that wild-type *P. pastoris* cells do not appreciably post-translationally modify heterologous actins either by *N*-acetylation or His-73 methylation, whereas Sf9 cells do. The ability to express unmodified actin, yet promote *N*-acetylation or His-73 methylation by expression of appropriate enzymes, as and when required, offers the potential to generate the correct actin isoform with the appropriate PTMs for biochemical, biophysical and structural studies. For example, study of *A. thaliana* actin *in vitro* could employ a mixture of Ala acetylated actin with a fraction (∼10%) that is His-73 methylated and the rest unmethylated. In this regard, we believe Pick-ya actin will facilitate the study of actins from a large variety of eukaryotes, including plants.

A simple flow chart for actin expression and purification requires establishing 1. if actin isolated from the source contains *N*-Met/Ala/Glu/Asp *N*-acetylation 2. if His-73 is methylated, and 3. if the *C*-terminal amino acid is a Phenylalanine. Based on the answers to these questions, expression of appropriate Ubiquitin-actin-Tβ4-8His fusion proteins with or without NAA80 and SETD3 and subsequent purification should provide pure actin isoforms with appropriate modifications.

## Materials and Methods

### Plasmids and strains

Plasmids and strains used in this study are listed in Table S1 and S2, respectively.

### pPICZc-actin-thymosin β4-8His

Actin-coding sequences codon-optimized for expression in *P. pastoris* were synthesized by gBlocks (IDT) and cloned into pPICZc (Invitrogen), which adds, at the *C*-terminus, an in-frame chymotrypsin cleavage site, linker sequence, thymosin β4 and a His tag (Noguchi et al., 2007; Hatano et al., 2018).

### *N*-terminal ubiquitin fusion constructs

Five’ part of *S. cerevisiae* UBI4 gene encoding ubiquitin repeat 1 (Ubi4 1-76 amino acid residues) were PCR amplified from genomic DNA isolated from *S.cerevisiae* and cloned at the 5’ end of in-frame actin in pPICZc-actin-thymosin β4-8His linearized by EcoRI using Gibson assembly. The first ATG for actin was removed by the cloning.

### NAA80 and SETD3 genes

NAA80 and *myc*-SETD3 genes codon-optimized for expression in *P. pastoris* were synthesised by gBlocks (IDT) and cloned into pIB2 (*PGAP* promoter) or pIB4 (*PAOX1* promoter) *P. pastoris* expression vectors by Gibson assembly.

### Dual expression of NAA80 and *myc*-SETD3

A PCR-amplified *PGAP*:NAA80∷*TAOX1* DNA fragment was cloned at the 3’ of NAA80 gene. The resulting plasmid lacking terminator for NAA80 gene was linealised by PstI/SphI digestion and *CYC1* terminator was cloned by ligation.

### *P. pastoris* culture

The composition of the minimal glycerol (MGY) and minimal methanol (MM) growth media and basic techniques for *P. pastoris* are described in the Pichia Expression Kit Instruction Manual (https://www.thermofisher.com/order/catalog/product/V19020).

### *P. pastoris* transformation

Plasmid DNA was linearized and yeast cells were transformed by using the lithium chloride method or electroporation. Transformants with pPICZc plasmid were selected on yeast extract peptone dextrose (YPD) solid media containing 100 mg/l Zeocin (Gibco, #R25001). For transformation of cells with pIB2 or pIB4 plasmids, *his4* cells were used as the host and transformants were selected on MGY. Single colonies were picked and cultured on YPD solid media and the transformants were re-selected to select stable clones.

### Purification of recombinant actin from *P. pastoris*

Actin was purified as described before with slight modifications. *P. pastoris* transformants were inoculated into MGY liquid medium and cultured at 30°C, with shaking at 220 rpm. The culture medium was diluted and cells were further cultured at 30°C, with rotation at 220 rpm, until the optical density at 600 nm (OD600) reached around 1 - 1.5. Cells were pelleted by centrifugation at 10628 g at 25°C for 10 min (Thermo Fisher Scientific, #F9-6×1000 LEX rotor) and washed once with sterilized water prior to re-suspension into (MM) medium. Cells were cultured in 2 l baffled-flasks (500 ml culture each) at 30°C, with rotation at 220 rpm, for 1.5–2 days; 0.5% methanol was added every 24 h of culturing. Cells were washed once with water and resuspended in water. The suspension was dripped into liquid nitrogen to freeze and stored at −80°C until use.

Frozen cells were loaded into a grinder tube (maximum 50 g of sample, #6801, SPEX® SamplePrep) pre-cooled with dry ice and ground in a Freezer mill (#6870, SPEX®SamplePrep) in a liquid nitrogen bath. The duration of the grinding was 1 min with 14 cycles per second (cps). The grinding was repeated 40 times at 1 min intervals. Liquid nitrogen was re-filled every 10 cycles of the grinding. The lysate powder was stored at −80°C until further use. The lysate was resolved in an equal amount of 2× binding buffer [20 mM imidazole, 20 mM HEPES pH 7.4, 600 mM NaCl, 4 mM MgCl_2_, 2 mM ATP, 2× concentration of protease inhibitor cocktail (cOmplete, EDTA free #05056489001, Roche), 1 mM phenylmethylsulfonyl fluoride (PMSF) and 7 mM β-mercaptoethanol (β-ME)]. The lysate was sonicated on ice (10 s with 60% amplitude, Qsonica Sonicators) until all aggregates were resolved. The lysate was centrifuged at 4°C at 3220 g for 15 min (Eppendorf #A-4-81 rotor) to remove intact cells and debris, then further cleared by centrifugation at 4°C and 25658 g for 1 h (Thermo Fisher Scientific, #A23-6×100 rotor). The supernatant was passed through a 0.22 µm filter (Corning #431097) and incubated with nickel resin (Thermo Scientific, #88222) at 4°C for 1–1.5 h. The resin was pelleted down by centrifugation at 4°C at 1258 g for 5 min (Eppendorf #A-4-81 rotor) and washed with 25 ml ice-cold binding buffer [10 mM imidazole, 10 mM HEPES pH 7.4, 300 mM NaCl, 2 mM MgCl_2_, 1 mM ATP, protease inhibitor cocktail (cOmplete, EDTA free #05056489001, Roche), 1 mM PMSF and 7 mM β-ME]. The resin was loaded into an open column (Bio-Rad, #731-1550) and washed first with ice-cold 20 ml binding buffer, then with 45 ml ice-cold G-buffer [5 mM HEPES (pH 7.4), 0.2 mM CaCl2, 0.01% (w/v) NaN_3_, 0.2 mM ATP and 0.5 mM dithiothreitol (DTT)]. The resin was resuspended in 6 ml ice-cold G-buffer containing 5 µg/ml TLCK-treated chymotrypsin (Sigma, #C3142) and incubated overnight at 4°C. The chymotrypsin was inactivated by addition of PMSF to 1 mM and incubated for 30 min on ice. The eluate was then collected into a tube. Actin retained on the resin was eluted with G-buffer and all the elution fractions were combined. The eluate was concentrated using a 30 kDa cut-off membrane (Sigma-Aldrich, #Z677892-24EA) and the final volume adjusted to 900 µl with ice-cold G-buffer. Actin was polymerized by addition of 100 µl 10× MEK solution [20 mM MgCl_2_, 50 mM glycol-bis(2-aminoethylether)-*N,N,N*′,*N*′-tetraacetic (EGTA) and 1 M KCl] for 1 h at room temperature. F-actin was pelleted by ultracentrifugation for 1 h at room temperature at 45,000 rpm (Beckman TLA-55 rotor) and re-suspended in ice cold G-buffer. F-actin was depolymerized by dialysis against 1 l G-buffer at 4°C for 2 days. Dialysis buffer was exchanged every morning and evening (total 3 times). Any remaining F-actin was pelleted by ultracentrifugation at room temperature at 45,000 rpm for 30 min (Beckman TLA-55 rotor). Then, actin in the supernatant was loaded into HiPrep 16/60 Sephacryl S-100 HR (GE healthcare, #17116501) pre-equilibrated with G-buffer for a size-exclusion chromatography (SEC). The concentration of actin was determined by measuring the absorbance at 290 nm [A290=0.63 (/mg/ml/cm)] using a NanoDrop 2000c spectrophotometer (Thermo Fisher Scientific).

### *S.cerevisiae* Act1 labelling with Alexa Fluor 488-maleimide or biotin-PEG2-maleimide

*S.cerevisiae* Act1 was expressed and purified as described above until digestion with chymotrypsin. After the digestion and inactivation of chymotrypsin by PMSF treatment, the eluate was collected and actin retained on the resin was also eluted with G-buffer without DTT. All the eluates were pooled in a tube and concentrated to 2.5 ml using a 30 kDa cut-off membrane (Sigma-Aldrich, #Z677892-24EA). The buffer was exchanged using a PD-10 desalting column with Sephadex G-25 resin (GE-Healthcare, #17085101) pre-equilibrated with G-buffer without DTT. The concentration of actin was adjusted to ∼25 µM and polymerized by addition of 100 mM KCl and 2 mM MgCl_2_. After 1 hour at 4°C, actin was incubated with 3-5 molar excess amount of Alexa Fluor 488 C5 maleimide (Invitrogen, #A10254) or EZ-Link Maleimide-PEG2-Biotin (Invitrogen, #A39261) for 1 hour at room temperature. The Cysteine-maleimide reaction was quenched by addition of 10 mM DTT to the sample and F-actin was pelleted, re-suspended in G-buffer as described above. Monomeric G-actin was obtained by deploymerisation and SEC as described above. The concentration of Alexa Fluor 488-labelled actin was measured by determining the absorbance at 290 nm (actin) and 495 nm (Alexa Fluor 488) on a NanoDrop 2000c spectrophotometer (Thermo Fisher Scientific), and calculated using the following formula: actin concentration (M)=A290/(0.63×42,000). The extinction coefficient of Alexa Fluor 488 at 495 nm is A495=72,000 M^−1^ cm^−1^, with a correction factor for Alexa Fluor 488 absorbance at 290 nm of 0.138. Labelling efficiency was calculated as follows: (A495/72,000)/actin concentration (M). The concentration of biotin-PEG2-labelled actin was calculated by SDS-PAGE and CBB staining using purified actin as the standard.

### Cover glass preparation for TIRF microscopy

Microscope cover glass (24 × 60 mm) was cleaned with 2% Helmanex II (Hellma, #9-307-011-4507) for 1 hour, followed by 3 M NaOH for 30 min in a 60°C water bath sonicator. Cleaned cover glass was rinsed with pure water and dried with nitrogen gas. The cover glass was adhering to sticky-Slide VI 0.4 (ibidi, #80608) to make 6 well flow chambers. The chamber was passivated with PLL-g-PEG/PEG-biotin mix [4 mg/ml PLL-g-PEG (,) and 0.05 mg/ml pll-peg-biotin (20%)] for 30 min at room temperature. The chamber was washed with G-buffer without ATP and DTT (G-AD buffer) prior to blocking with 1% BSA in G-AD buffer. The chamber was then washed with G-AD buffer prior to incubation with 0.1 mg/ml avidin in G-AD buffer for 15 min at room temperature. The chamber was washed with G-AD buffer, followed by TIRF-buffer [10 mM imidazole (pH 7.4), 50 mM KCl, 1 mM MgCl_2_, 1 mM EGTA, 0.2 mM ATP (pH 7), 10 mM DTT, 0.5% methylcellurose (4000 cP) and 15 mM glucose].

### Actin polymerization assay with TIRF microscopy

20 µM actin consists of ∼80% target actin, 20% Alexa Fluor 488-Act1 and 0.25% biotin-PEG2-Act1 were ultracentrifuged at ∼25 psi with Airfuge for 10 min at room temperature. The top 75% supernatant fraction was collected as the sample. Actin was diluted to 1.5 µM in TIRF buffer containing 12 kU/ml catalase and 48 kU/ml glucose oxidase to start its polymerization.

### TIRF microscopy settings and image acquisition

Andor Revolution TIRF system was equipped with the inverted Nikon Eclipse microscope base, Nikon Apo 100x/1.49 NA Apo TIRF objective lens, Nikon TIRF module, a 60 mW 488 nm solid state Laser and an Andor Zyla sCMOS camera. Images were acquired at 65 nm/pixel. The TIRF microscopy was done at room temperature. Images were acquired with 2 second interval.

### Mass spectrometric analysis

The protein was reduced using 10 mM DTT (#MB1015, Melford) for 60 min at room temperature. The sample was then alkylated with 55 mM iodoacetamide (#I6709, Sigma) for 20 min in the dark at room temperature and digested using 1 μg trypsin (sequencing grade; #V5111, Promega) per 100 μg of protein overnight at 37°C. Twenty microlitre samples were then analysed by nano LC-ESI-MS/MS using UltiMate® 3000 HPLC series using Nano Series™ Standard Columns for separation.A linear gradient from 4% to 25% solvent B (0.1% formic acid in acetonitrile) was applied over 30 min, followed by a step change 25–35% solvent B for 10 min, followed by a 3 min wash of 90% solvent B. Peptides were directly eluted (∼250nl/min) via a Triversa Nanomate nanospray source into a Orbitrap Fusion mass spectrometer (Thermo Scientific). Positive ion survey scans of peptide precursors from 375 to 1575 m/z were performed at 120 K resolution (at 200m/z) with automatic gain control 2 × 105. Precursor ions with charge state 2–6 were isolated and subjected to HCD fragmentation in the IonTrap at 120 K. MS/MS analysis was performed using collision energy of 33%, automatic gain control 5 × 103 and max injection time of 150 ms. The dynamic exclusion duration was set to 25 s with a 10 ppm tolerance for the selected precursor and its isotopes. Monoisotopic precursor selection was turned on. Label-free quantification (LFQ) of modified precursor ions and unmodified peptides was performed using MaxQuant (V1.5.5.1). The LFQ module calculates the integrated peak area of the precursor ion based on a retention time window of 5 min and ppm error window of 10 ppm to quantify the presence and absence of a precursor ion of interest in different mass spectrometry output files. LFQ intensity data for each sample were normalized to top 3 unmodified actin peptides. Data was analyzed in triplicates.

## Acknowledgements

We wish to thank members of the Balasubramanian laboratory, especially Dr. Saravanan Palani and Dr. Luke Springall for discussion and critical comments. We thank Dr. Suresh Subramani for *P.pastoris* plasmids and strains. We thank Dr. Rob Cross, who coined the Pick-ya actin acronym. We thank the Warwick Proteomics RTP for mass spectrometry.

## Author contributions

Conceptualization: T.H., M.K.B.; Methodology: T.H.; Validation: T.H., L.S., H.H.; A.S.; Formal analysis: T.H., L.S., H.H.; Investigation: T.H., L.S., H.H., A.S.; Resources: T.H., M.K.B.; Writing -original draft: T.H., M.K.B.; Writing -review & editing: T.H., L.S., H.H.; A.S., M.K.B.; Supervision: M.K.B.; Project administration: T.H., M.K.B.; Funding acquisition: M.K.B.

